# Alternative splicing is a developmental switch for hTERT expression

**DOI:** 10.1101/2020.04.02.022087

**Authors:** Alex Penev, Andrew Bazley, Michael Shen, Jef D. Boeke, Sharon A. Savage, Agnel Sfeir

## Abstract

High telomerase activity is restricted to the blastocyst stage of embryonic development when telomere length is reset, and is characteristic of embryonic stem cells (ESCs) and induced pluripotent stem cells (iPSCs). However, the pathways involved in telomerase regulation as a function of pluripotency remain unknown. To explore hTERT transcriptional control, we compare genome-wide interactions (4C-seq) and chromatin accessibility (ATAC-seq) between human ESCs and epithelial cells and identify several putative hTERT *cis-*regulatory elements. CRISPR/Cas9-mediated deletion of candidate elements in ESCs reduces the levels of hTERT mRNA but does not abolish telomerase expression, thus implicating post-transcriptional processes in telomerase regulation. In agreement with this hypothesis, we find an hTERT splice variant lacking exon-2 and prone to degradation, to be enriched in differentiated cells but absent from ESCs. In addition, we show that forced retention of exon-2 prevents telomerase silencing during differentiation. Lastly, we highlight a role for the splicing co-factor SON in hTERT exon-2 inclusion and identify a SON mutation in a Dyskeratosis congenita patient with short telomeres and decreased telomerase activity. Altogether, our data uncover a novel alternative splice switch that is critical for telomerase activity during development.

---

Telomerase counteracts telomere erosion and promotes cellular immortality. This specialized ribonucleoprotein complex is minimally composed of a reverse-transcriptase protein subunit (TERT) and an integral telomerase RNA component (TR). While the expression of telomerase RNA is ubiquitous (1, 2), TERT levels are tightly regulated (3). Human TERT (hTERT) is turned on during the blastocyst stage of embryonic development when telomere length is re-established (4), and subsequently downregulated in most somatic cells (4, 5). Similarly, hTERT becomes activated during the process of nuclear reprogramming (6) and is essential for telomere maintenance in pluripotent cells (7). Upregulation of telomerase occurs in approximately 90% of cancers, allowing tumor cells to escape crisis and proliferate indefinitely (5, 8). Somatic mutations in the hTERT promoter have been reported in a subset of cancers and shown to increase telomerase activity by creating a binding site for ETS family transcription factors (9-11). To date, the underlying mechanism that activates hTERT during development and its subsequent silencing in somatic tissues remains unknown.

The core hTERT promoter contains numerous binding sites for pluripotency and growth-related transcription factors, including Myc, Klf4, and Sp1, but expression of these different factors is not sufficient for hTERT accumulation and telomerase activation in somatic cells (12). We therefore sought to identify *cis*-regulatory elements (REs) that could account for the difference in hTERT mRNA levels between pluripotent and somatic cells (Figure S1A). Active REs are known to directly contact the promoters they regulate and have increased chromatin accessibility, we therefore, performed <u>C</u>ircular <u>C</u>hromosomal <u>C</u>onformation <u>C</u>apture (4C) and ATAC-seq in H7 human embryonic stem cells (ESC) and adult retinal pigment epithelial (ARPE) cells that are proficient for p53 and have a stable diploid karyotype. We used a 1.1 kb fragment spanning the hTERT promoter as bait and captured sequences that are in close proximity by 4C (13), using 4Cker software to analyze Illumina sequencing reads (Figure S1B-C) (14). Our analysis identified loci that contact the hTERT promoter exclusively in telomerase positive cells (H7) or in telomerase negative cells (ARPE), as well as loci that contact the TERT promoter in both cases. Notably, the majority of 4C-interactions occurred within the boundaries of the topologically associated domain (TAD) within which the hTERT locus resides (Figure 1A and Figure S1C). In parallel, we profiled genome-wide chromatin accessibility in H7 and ARPE cells using ATAC-seq (15). As expected, ATAC-seq revealed regions of open chromatin specific to each cell type, including the hTERT promoter in ESCs (Figure S1D).

**Figure. 1.**
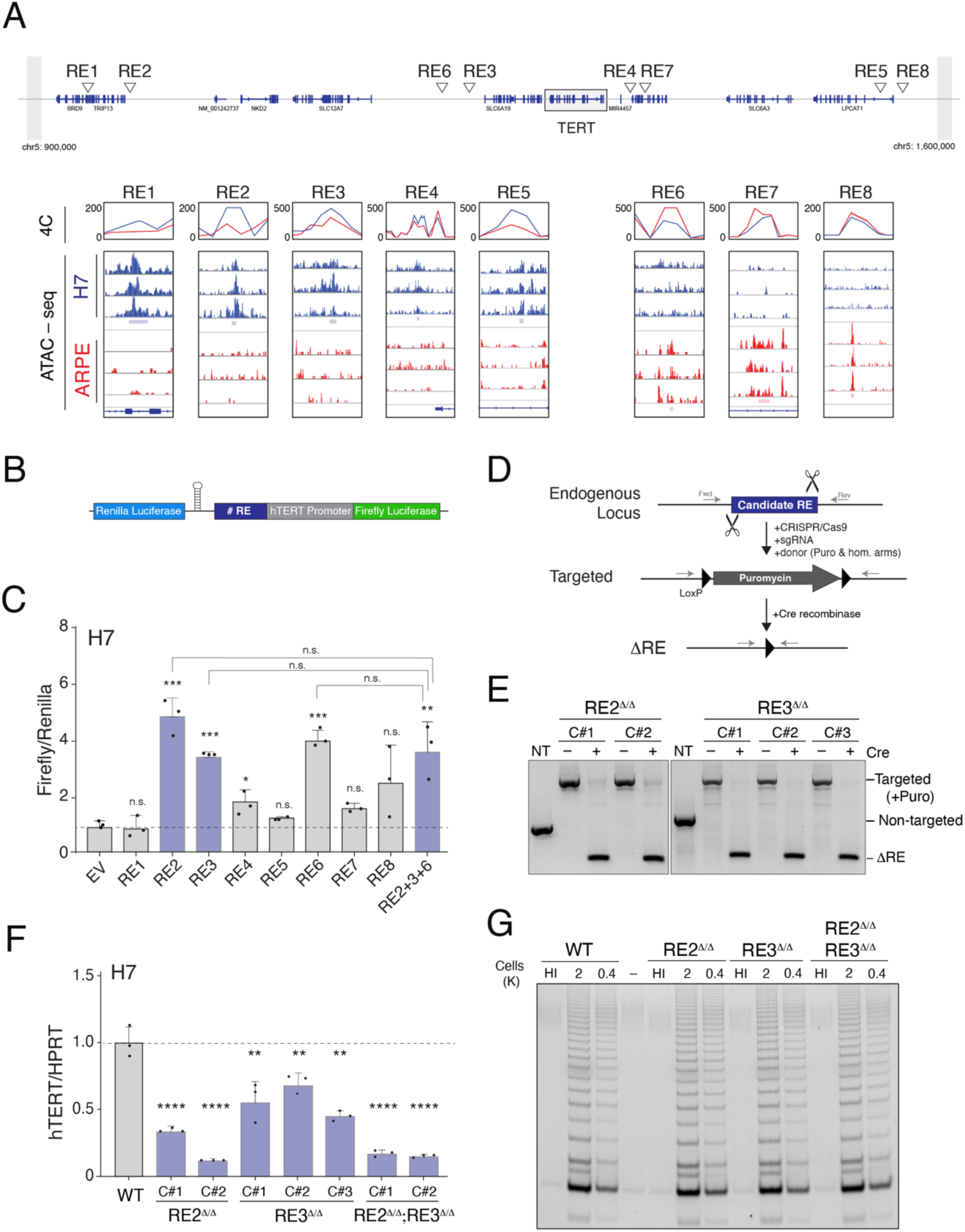
Identification of novel cis-regulatory elements (REs) that modulate hTERT expression as a function of pluripotency. A) Top: Refseq of annotated genes on chr5, depicting candidate RE elements within the TAD boundary surrounding the hTERT locus (grey box). Bottom: Overlay of data obtained from ATAC-seq and 4C-seq analysis to identify candidate hTERT REs. Candidate enhancers (left, RE1-RE5) preferentially interact with hTERT promoter and display open chromatin in embryonic stem cells (ESCs). Silencer elements (right, RE6-RE8) interact with hTERT promoter in differentiated cells. 4Cker software was used to determine interaction score (line graphs) and was aligned to ATAC-seq results from three independent replicates. B) Schematic of the dual luciferase reporter used to interrogate the function of individual REs. hTERT core promoter is composed of 500bp sequence upstream of transcription start site (TSS) as well as 100bp downstream sequence encompassing hTERT start codon. Individual REs were cloned upstream of hTERT promoter. Expression of Renilla luciferase driven by a constitutive promoter for normalization. A strong hairpin is positioned downstream of the Renilla gene to prevent transcriptional read-through. C) Bar graph represents the ratio of Firefly to Renilla luciferase in ESCs expressing the reporter plasmid with the indicated RE (n=3, *: p<0.05, **: p<0.01, ***: p<0.001). D) Schematic of CRISPR/Cas mediated gene editing of RE2 and RE3 in human ESCs. Cells were co-transfected with two sgRNA that cut on either side of the 500bp locus and a double-stranded (ds) DNA donor containing Puromycin resistance cassette. Following targeting, excision of Puromycin cassette was carried out using lentiviral Cre recombinase. Arrows indicate primers used for genotyping. E) Genotyping PCR on cells with the indicated genotype and Cre-treatment using primers highlighted in (D). PCR products (RE2/RE3): wild type, 560/700bp; targeted, 1500/1700bp and null alleles, 250/300bp F) Quantitative RT-PCR for hTERT mRNA in cells with the indicated genotype following treatment with Cre recombinase. (n=3, RE3^Δ*/*Δ^: p<0.01, RE2^Δ*/*Δ^ and DKO: p<0.0001). G) Telomere Repeat Addition Processivity (TRAP) assay to measure telomerase holoenzyme activity in clonally derived ESCs lacking the indicated RE. HI=heat inactivated

By superimposing 4C interactions onto the chromatin accessibility by ATAC-seq, we identified a number of regions that displayed increased hTERT promoter interaction and open chromatin and are specific to either H7 or ARPE (Figure 1A). We identified five putative hTERT enhancers elements, termed Regulatory Elements (RE) 1-5, that displayed increased contact with hTERT promoter in H7 cells and showed an open chromatin configuration (Figure 1A). With the exception of RE4, which was reported to enhance hTERT activity in cancer cell lines, these potential enhancer elements have not been previously detected (16). Our analysis also recognized three putative hTERT transcriptional silencers elements (RE6-RE8) that show enhanced interaction with the hTERT promoter and increased chromatin accessibility in differentiated cells (Figure 1A).

To validate the putative REs, we introduced sequences corresponding to RE1-RE8 upstream of hTERT core promoter in a dual-luciferase reporter plasmid and assayed for luciferase activity in H7 and ARPE cells (Figure 1B and S1E-F). Among those tested, RE2 and RE3 significantly enhanced hTERT promoter activity in ESCs but not in somatic cells (Figure 1C and S1F). Paradoxically, the putative repressor element, RE6, also showed enhanced promoter activity in H7 cells. The latter can be explained if RE6 contained a transcription factor binding site that is only accessible when removed from its genomic context. We combined RE2, RE3, and RE6, tandemly in the luciferase reporter and detected no additive effect on hTERT promoter activity (Figure 1C). The latter observations potentially rule out cooperativity in enhancer function during hTERT transcription. To further characterize the putative regulatory elements, we mined publicly-available ENCODE ChIP-seq data for histone modifications that distinguish active enhancers (17). Consistent with RE2 and RE3 acting as enhancers, we detected significant deposition of H3K4me1 at these loci, whereas no enrichment of H3K4me1 was evident at the RE6 locus (Figure S1G).

We next tested the enhancer function of RE2 and RE3 *in vivo* using CRISPR/Cas9 genome editing of ESCs. Our strategy entailed replacing individual enhancers with a donor sequence containing a Puromycin resistance cassette and flanked by two LoxP sites (Figure 1D). We derived independent ESC clones for each locus and confirmed homozygous targeting by genotyping PCR and Sanger sequencing (Figure 1E and data not shown). Targeted cells were then transduced with a lentiviral-encoded Cre-recombinase to excise the puromycin cassette yielding RE2^Δ*/*Δ^ and RE3^Δ*/*Δ^ cells (Figure 1E). We performed quantitative RT-PCR and observed two to four-fold reduction in hTERT mRNA levels in enhancer-deleted cells, confirming enhancer activity for both RE2 and RE3 (Figure 1F). In addition, we generated double-knockout RE2^Δ*/*Δ^RE3^Δ*/*Δ^ ESCs and observed that dual targeting of both enhancers did not lead to further reduction in hTERT transcript when compared to RE2^Δ / Δ^ cells (Figure 1F and Figure S1H). These results provided further support that the two enhancers do not act cooperatively during hTERT transcriptional activation. Despite the noted reduction in hTERT mRNA levels, Telomere Repeat Addition Processivity (TRAP) assay revealed that cells lacking RE2 and RE3 had similar telomerase activity as non-targeted cells (18) (Figure 1G). Given that the reverse transcriptase encoded by hTERT is not rate limiting for telomerase activity in human ESCs (11), a four-fold reduction in hTERT mRNA is not expected to impact overall telomerase activity in H7 cells. In summary, our analysis identified RE2 and RE3 as novel enhancer elements that modulate hTERT mRNA levels in ESCs *vs.* differentiated cells. Nonetheless, the maximal reduction for telomerase transcripts in enhancer-deleted cells was significantly less than the observed 100-fold difference in hTERT expression between pluripotent and differentiated cells (Figure S1A). While we cannot rule out that additional distant enhancer elements might further regulate hTERT transcription, it is also possible that transcriptional regulation alone does not account for the strong induction of telomerase in ESCs and its subsequent repression in differentiated cells. To test this idea, we transduced wildtype fibroblasts with Vp64-dCas9 fusion protein and two guide RNAs complementary to the hTERT promoter (19). We then performed RT-qPCR and observed minimal hTERT activation in Vp64-dCas9 expressing fibroblasts relative to ESCs (Figure S1I). Based on these results, we concluded that recruitment of a robust transcriptional activator to the hTERT promoter in differentiated fibroblasts led to only mild accumulation of telomerase mRNA.

We next sought to explore the impact of post-transcriptional processing on the abundance and stability of hTERT mRNA in ESCs. Specifically, we focused on alternative splicing of pre-mRNA as a regulatory pathway that has been previously shown to control developmentally-regulated genes and pluripotency-associated factors (20-25). Many hTERT alternative splice variants have been detected in different cell types, including α/β variants that modulate telomerase activity in cancer cells (26-28). We analyzed the full spectrum of telomerase splice variants in H7 and ARPE cells using RNA Capture-seq that combines tiling arrays of biotinylated oligos with RNA-seq and therefore, enhances the sequencing coverage for low abundance transcripts such as hTERT. hTERT Capture-seq identified notable reads spanning exon-1 and exon-3, indicative of hTERT transcripts lacking exon-2 (hTERT-ΔEx2), that were abundant in differentiated cells but not ESCs (Figure 2A-B). These results are consistent with a previous meta-analysis, in which transcripts lacking exon-2 were consistently and widely expressed in many human cell types except ESCs (29).

**Figure. 2.**
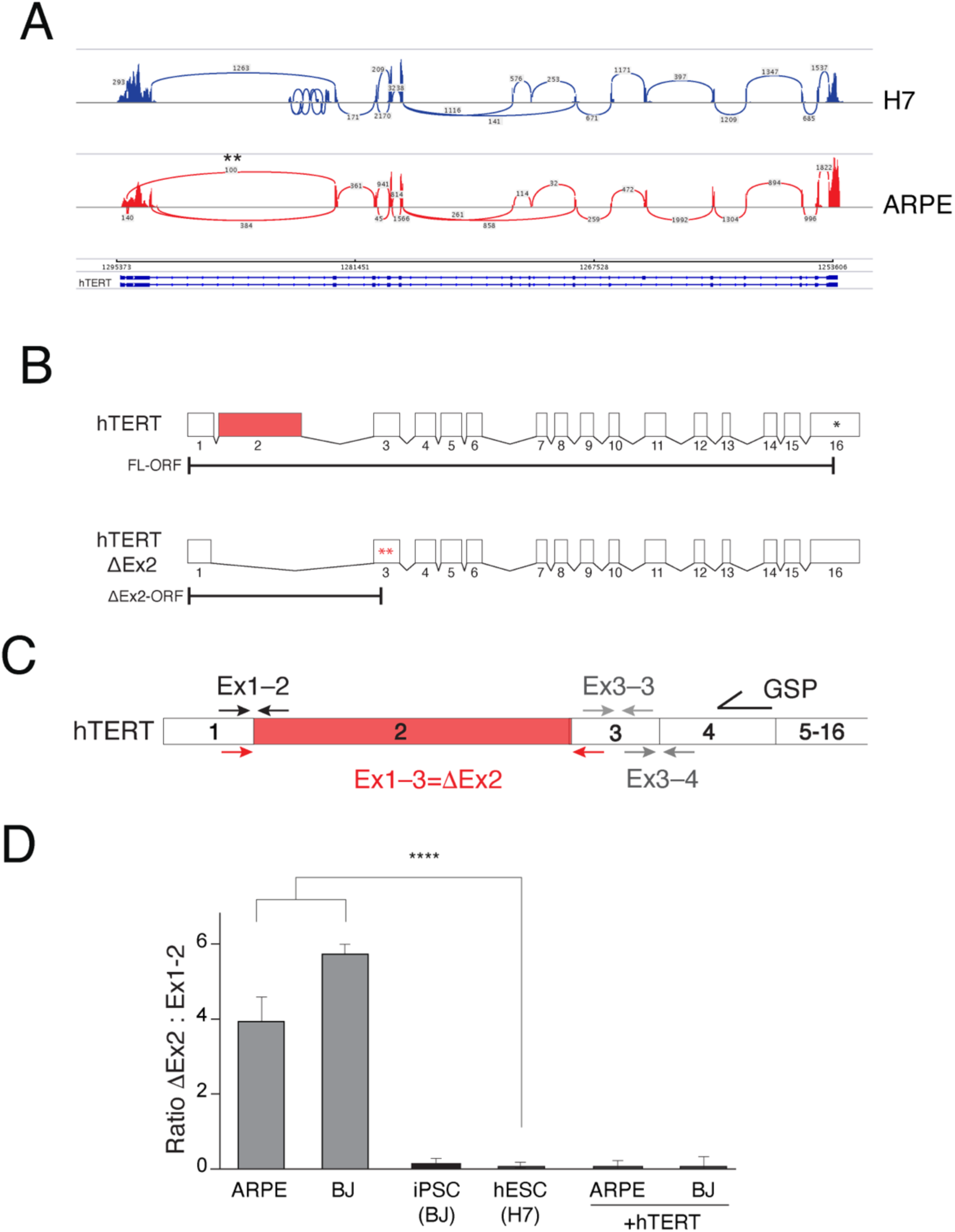
Inclusion of exon-2 correlates with the abundance of telomerase mRNA. A) Sashimi plot representing RNA Capture-Seq for hTERT locus in H7 ESCs and ARPE cells. Reads from exons are depicted as pileups and exon-exon junctions denoted with arcs. Asterisks indicate ΔEx2 splice variant transcripts detected in differentiated cells. B) Schematic of full-length and hTERT-ΔEx2 transcripts, demonstrating the position of the premature termination codons (PTC, asterisks) generated upon exon-2 skipping. Two tandem PTC’s would likely target ΔEx2 transcript for degradation by nonsense-mediated decay (NMD). C) Schematic illustration of junction-spanning PCR strategy used to assay hTERT ΔEx2 abundance by quantitative RT-PCR. RNA was reverse-transcribed using an hTERT gene-specific primer (GSP) and cDNA was purified and equalized between samples prior to PCR amplification with the indicated primers. D) Quantification of the ratio of hTERT ΔEx2 relative to full-length as determined by qRT-PCR in mortal cell lines (ARPE & BJ), ARPE and BJ cell lines immortalized with hTERT cDNA, iPSCs derived from BJ cells, and human ESCs (n=3, p<0.0001).

To corroborate the results of the RNA Capture-Seq, we designed a junction-specific PCR strategy in which cDNA was first generated with a gene-specific primer in hTERT exon-4 and subjected to exon-spanning quantitative PCR to determine the ratio of hTERT-ΔEx2 relative to full-length hTERT mRNA (Figure 2B-C). Our analysis revealed that hTERT-ΔEx2 was 4 times more abundant than full-length transcript in cells lacking telomerase activity, including epithelial cells (ARPE) and fibroblasts (BJ). In contrast, ESCs and iPSCs derived from BJ fibroblasts displayed diminished levels of hTERT-ΔEx2. In conclusion, junction-PCR analysis further confirmed a strong correlation between exon-2 skipping and the absence of telomerase activity in differentiated cells (Figure 2D). Importantly, we detected the same correlation between telomerase activity and exon-2 alternative splicing in RNA samples obtained from human tissues, including brain (telomerase negative) and testis (telomerase positive) (Figure S2A).

Exclusion of hTERT exon-2 is predicted to generate tandem premature termination codons in exon-3 that would act as signal for mRNA decay (30) (Figure 2B). We examined the effect of inhibiting the RNA decay machinery on the abundance of hTERT transcripts. To that end, we performed junction-specific RT-qPCR following the transient depletion of nonsense-mediated decay factor, UPF1. Our results revealed increased accumulation of hTERT-ΔEx2 transcripts in HeLa cells lacking UPF1 (Figure S2B). We obtained similar results upon treatment of cells with small molecule inhibitors for the RNA decay machinery (Figure S2C), suggesting that exon-2 skipping yields transcripts that are susceptible to degradation. In a complementary approach, we transfected HeLa cells with splice-blocking anti-sense morpholinos (ASOs) that specifically prevented the inclusion of exon-2. This then led to increased accumulation of hTERT-ΔEx2 variant mRNA and a concomitant reduction in transcripts containing exon-2 (Figure S2D-E). In agreement with exon-2 exclusion compromising telomerase function, we noted a significant reduction in TRAP activity in cells treated with exon-2 ASO relative to control cells (Figure S2F-G). Taken together, our results indicate that selective transcripts lacking exon-2 trigger the nonsense-mediated decay machinery to degrade hTERT mRNA and compromise telomerase function.

So far, our data uncovered a strong correlation between exon-2 inclusion and the accumulation of hTERT transcripts. We next tested the impact of forced exon-2 retention on telomerase silencing during differentiation by targeting the first intron of hTERT in human ESCs with CRISPR/Cas9 (Figure 3A). In effect, deletion of hTERT intron-1 results in a constitutive fusion between exon-1 and exon-2 and thus prohibits the alternative splicing event that results in exon-2 exclusion. We performed two separate rounds of gene-editing and isolated a total of six independent hTERT^Δin1/Δin1^ clones that we validated using genotyping PCR and Sanger sequencing (Figure 3B and S3A). hTERT^Δin1/Δin1^ ESCs maintained proper morphology and expressed appropriate cell surface markers (Figure S3B) (6). Furthermore, telomerase activity in hTERT^Δin1/Δin1^ ESCs was similar to that of hTERT^+/+^ cells (Figure S3D). We then differentiated hTERT^Δin1/Δin1^ ESCs into fibroblasts and confirmed downregulation of key pluripotency genes and a concomitant upregulation of fibroblast-specific genes (Figure S3B-C). As expected, we observed complete repression of hTERT upon differentiation of hTERT^+/+^ ESCs (Figure 3C) (4, 31). In contrast, hTERT^Δin1/Δin1^ fibroblasts retained significant hTERT mRNA levels by quantitative RT-PCR (Figure 3C). To further corroborate the failure of hTERT silencing in differentiated hTERT^Δin1/Δin1^ cells, we quantified hTERT mRNA directly using custom NanoString™ probes and noted the retention of sequences corresponding to several exons (Figure 3D). Additionally, we examined hTERT mRNA abundance and localization using single-molecule RNA-FISH probes (sm-FISH) (32) and confirmed its accumulation in hTERT^Δin1/Δin1^ but not hTERT^+/+^ fibroblasts (Figure S3E). sm-FISH revealed that hTERT transcripts in hTERT^Δin1/Δin1^ cells are capable of being exported from the nucleus, albeit not as efficiently as pre-spliced hTERT cDNA. Lastly, we showed that the abrogation of hTERT silencing was not limited to hTERT^Δin1/Δin1^ fibroblasts but was also evident when ESCs were differentiated into hepatocyte-like cells *in vitro* (Figure S4A-B) (33).

**Figure 3.**
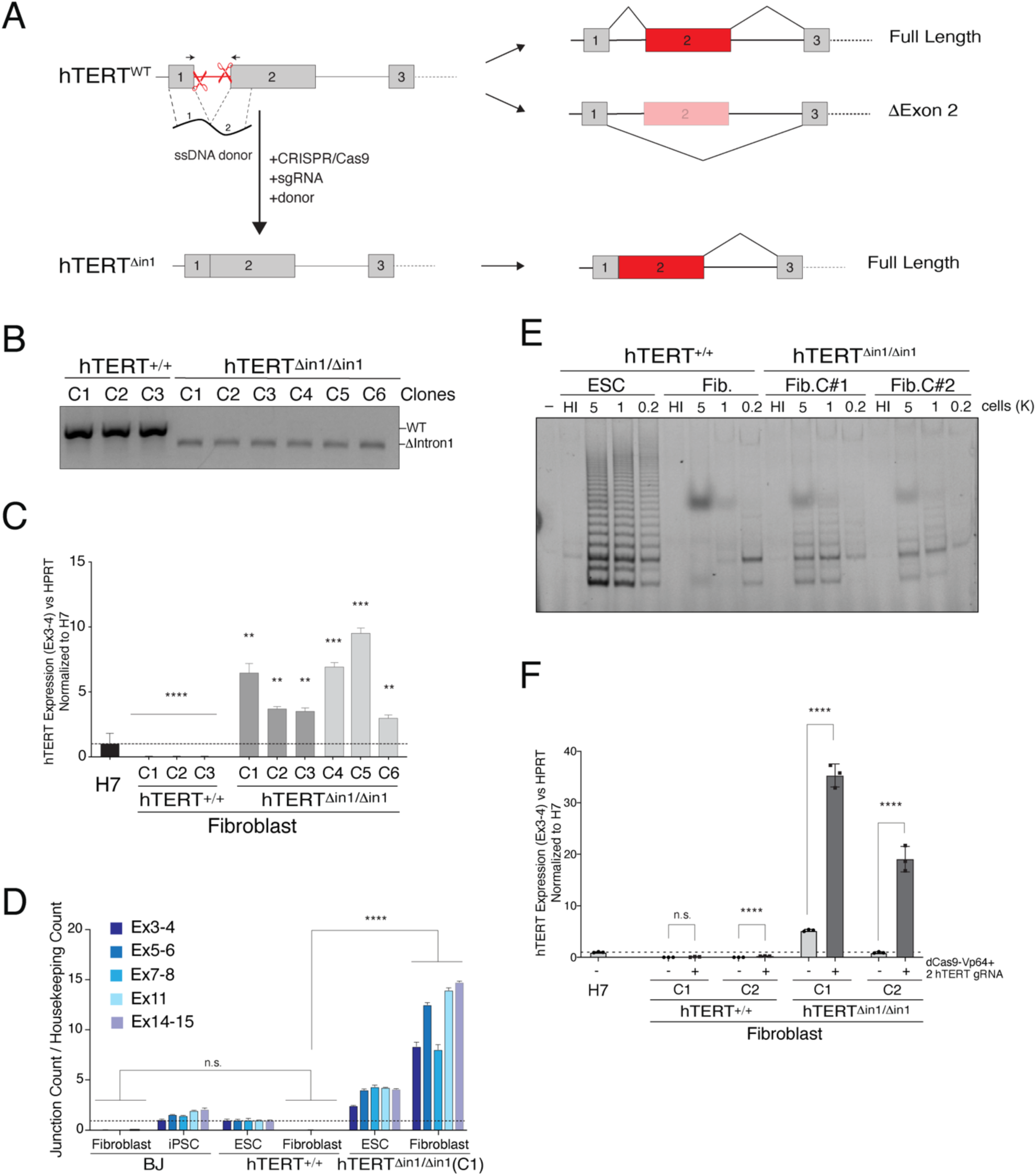
Forced retention of exon-2 abolishes silencing of hTERT upon differentiation. A) Schematic illustration of intron 1 deletion by CRISPR/Cas9 gene editing and the predicted splicing pattern. Cells were co-transfected with two sgRNA that cleave within hTERT intron-1 and a 200bp single-stranded (ss) DNA donor containing 100bp sequence from exon-1 and exon-2 directly concatenated. B) Genotyping PCR from cells with the indicated genotype. PCR products: wild type, 584bp; Δintron1, 480bp C) Quantitative RT-PCR for hTERT mRNA in cells with the indicated genotype. hTERT expression is silenced in fibroblasts derived from wildtype ESC clones, whereas hTERT levels remain elevated upon differentiation of hTERT^Δin1/Δin1^ ESCs. Values are normalized to hTERT^+/+^ H7 ESC (n=3, **: p<0.01, ***: p<0.001, ****: p<0.0001). D) Absolute quantification of multiple hTERT exon-exon junctions using direct Nanostring quantification of RNA confirms increased hTERT transcripts in fibroblasts derived from hTERT^Δin1/Δin1^ ESCs. Data normalized to hTERT^+/+^ ESCs (n=3, p<0.001). E) TRAP assay for telomerase activity in cells with the indicated genotype shows that hTERT^Δin1/ Δin1^ fibroblasts retain telomerase activity compared to wildtype control cells. F) RT-qPCR for hTERT mRNA in differentiated fibroblasts in the context of Vp64-Cas9 transcriptional activation of hTERT promoter. Two hTERT ^+/+^ and hTERT^Δin1/Δin1^ clones were transduced with lentivirus encoding Vp64-dCas9 transactivation protein and 2 guide RNAs complementary to hTERT promoter. Following FACS selection of Vp64-dCas9 expressing cells, RT-qPCR was performed on non-transduced parental clonal lines and Vp64-dCas9 expressing cells. Values normalized to H7 cells (n=3, p<0.01).

To rule out that intron-1 contained a cis-regulatory element that enforced hTERT silencing in differentiated cells, we introduced the corresponding sequence into the dual-luciferase reporter assay and detected no difference in hTERT promoter activity (Figure S4C). Additionally, we generated clonally-derived ESCs in which hTERT intron-1 was replaced with scrambled intronic sequence that retained the splice junctions (Figure S4D), and found that hTERT mRNA levels were similar to non-targeted ESCs (Figure S4E). In conclusion, our results demonstrated that forced retention of hTERT exon-2 prevents telomerase silencing in differentiated cells. We then examined telomerase enzymatic function and observed increased TRAP activity in hTERT^Δin1/Δin1^ fibroblasts relative to differentiated hTERT^+/+^ cells (Figure 3E and Figure S4F). These results further support that exclusion of exon-2 is a key step during telomerase silencing upon differentiation. It is worth noting that although hTERT^Δin1/ Δin1^ fibroblasts retained significant levels of hTERT mRNA, telomerase activity in the targeted fibroblasts was significantly lower than ESCs. It is established that modification of pre-mRNA influences post-transcriptional processes (34, 35), including mRNA modification, nuclear export, and translation efficiency that could be compromised in differentiated cells.

Having explored hTERT regulation by transcriptional activation and alternative splicing separately, we next sought to investigate the interplay between the two processes during telomerase activation. To that end, we exploited the dCas9-Vp64 transactivation system to increase hTERT promoter activity in hTERT^Δin1/ Δin1^ and hTERT^+/+^ fibroblasts, independently (Figure 3F and S4G). Analysis of hTERT mRNA by quantitative RT-qPCR revealed that while transcriptional activation has\d a minimal impact in wildtype cells, a synergistic effect on hTERT mRNA accumulation was observed when Vp64-dCas9 was recruited to the hTERT promoter in hTERT^Δin1/ Δin1^ cells (Figure 3F). In summary, we conclude that transcriptional activation is not sufficient to mount robust hTERT levels in differentiated cells. Instead, our data is consistent with promoter activation being coupled to exon-2 retention by alternative splicing to allow the accumulation of hTERT mRNA.

In order to provide insight into the mechanism underlying alternative splicing of hTERT exon-2 and identify factors that influence exon choice, we carried out a small-scale RNAi-screen using luciferase-reporter genes. We generated two synthetic doxycycline-inducible “minigenes” comprising hTERT exons 1 to 3 and the spanning introns (Figure 4A). In the first minigene, Firefly luciferase was in-frame with the hTERT ORF containing exon-2 and therefore expressed only when the second exon was incorporated into the mRNA. Conversely, in the second minigene, Nano-luciferase was placed downstream of hTERT-ΔEx2 reading frame and was specifically expressed upon exclusion of exon-2. Both minigenes were assembled in yeast by homologous recombination (36) and included FRT sites to facilitate their integration into T-REx™-HeLa cells (Figure S5A-B). HeLa cells express hTERT-ΔEx2 and full-length transcripts at similar levels and were therefore suitable to investigate factors that regulate telomerase alternative splicing (Figure S5C). We co-transfected HeLa-FRT cells with minigenes I & II and established a clonally derived cell line with heterozygous integration of both minigenes at the same locus. In order to calculate the ratio of hTERT-ΔEx2 transcripts relative to full-length mRNA, we measured Firefly and Nano luciferase activity 48 hours after doxycycline treatment (Figure S5D). Upon validating the minigene reporter line, we assembled an siRNA mini-library targeting 442 genes annotated as RNA-binding proteins and splicing factors (Supp. Table 3) (37) and performed a small-scale RNAi screen. We monitored dual luciferase activity and found that knockdown of 77 genes significantly altered Nano:Firefly ratio (p<0.05) (Figure 4B). Top siRNA hits that enhanced inclusion of exon-2 included RBM14, a paraspeckle protein involved in nuclear RNA sequestration (38) and Mbnl1, a splicing regulator that represses pluripotency-associated exon inclusion (23). On the other hand, candidates that increased the ratio of Nano:Firefly luciferase and therefore, promoted the exclusion of hTERT exon-2 comprised multiple SF3A/B family members and the testis specific factor, BrdT.

**Figure 4.**
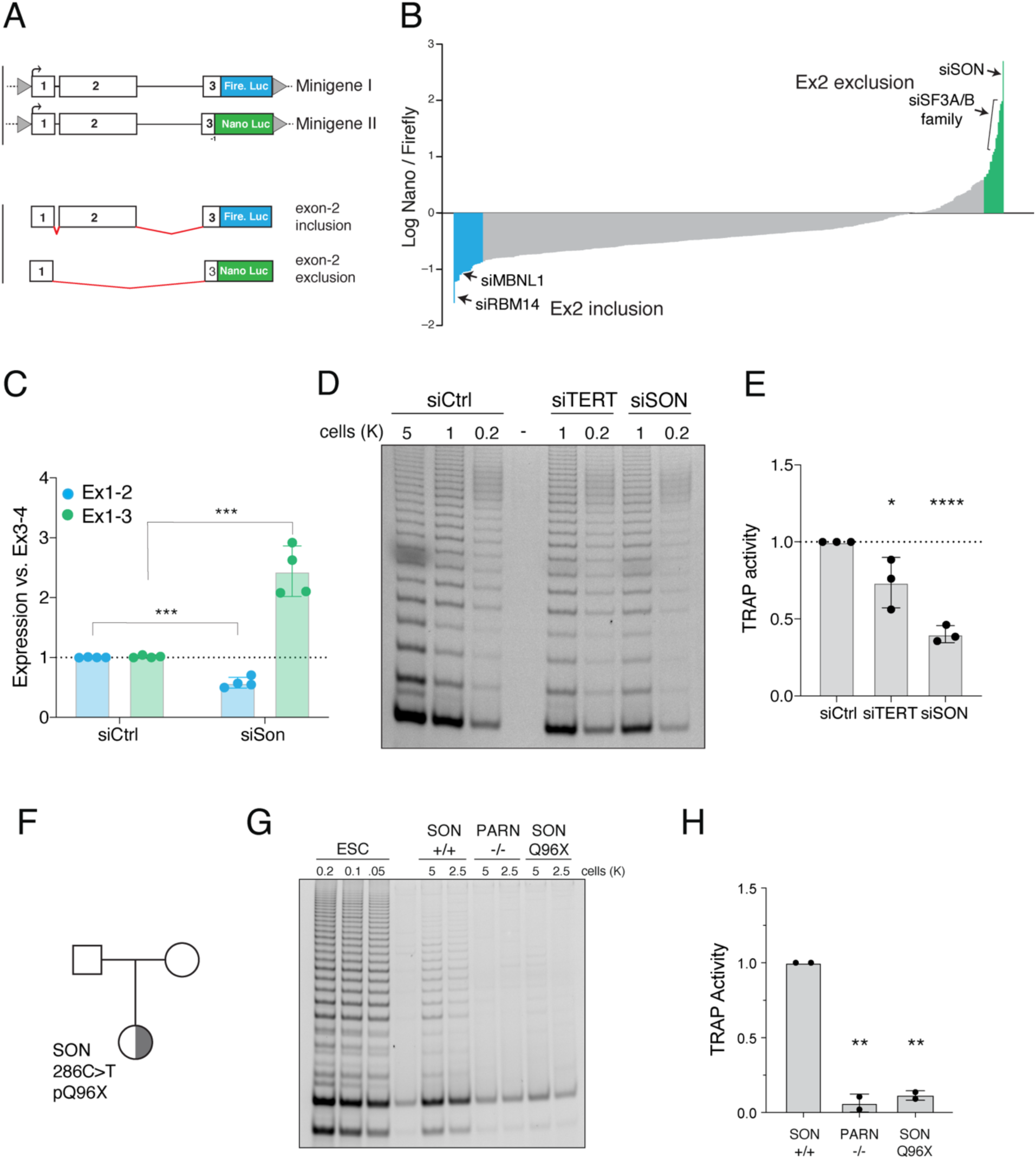
SON is a key regulator of hTERT alternative splicing. A) Schematic of minigenes designed to measure the efficiency of hTERT mRNA splicing using Nano- and Firefly luciferase activity as a readout. Retention of exon-2 leads to expression of Firefly luciferase, but not Nano luciferase. Conversely, exclusion of exon 2 leads to a (−1) frameshift in exon-3 prompting the expression of Nano-luciferase while shifting Firefly luciferase out-of-frame. B) The minigenes were integrated into HeLa cells and a small-scale RNAi screen using a curated list of splicing factors and RNA-binding proteins was performed. Graph depicts the average ratio of Nano/Firefly luciferase of 3 biological replicates for 442 genes. Data presented as a log-ratio, colors highlight genes with p-value <0.05. Genes highlighted in green are putative positive regulators of hTERT, and genes in blue are negative regulators of hTERT expression. C) Quantification of the abundance of hTERT exon1-2 and exon1-3 splice-junctions in ESCs following knockdown of SON. (n=4, p<0.001). D) Representative TRAP assay to detect telomerase activity in ESCs, 48 hours after SON depletion with siRNA. E) Quantification of the TRAP assay as in D. F) Pedigree highlighting the proband, a female child of unaffected parents, carrying a de novo SON Q96X heterozygous mutation (Q96X) affected with ZTTK and DC. G) Representative TRAP assay for telomerase activity in PBMC from a healthy donor (WT) and the proband (SON). PBMC from a PARN patient was used as a control H) Quantification of the TRAP assay in (G) (p<0.01). WT – TA 4646 0523. PARN – NCI-382-1 – TA 2812 0860 mutation p.N7H (c.19A>C) and deletion chr16:14,037,911 - 15,319,123 (patient published in (45)). SON – NCI-550-1– TA 5330 0534 mutation c.286C>T exon 3 p.Q96X

The strongest hit from our small-scale RNAi screen was the nuclear speckle protein and alternative splicing co-factor, SON. Knockdown of SON resulted in a significant increase in the ratio of Nano:Firefly luciferase, indicative of strong exon-2 exclusion. SON is enriched in ESCs and some cancer cells and has been shown to be required for splicing of pluripotency genes by facilitating the inclusion of exons with weak consensus splicing sites (39) (40). Given that long-term inhibition of SON was previously shown to impair pluripotency (39), we transiently depleted the gene in H7 and HeLa cells using siRNA (Figure S5E). As a control, we showed that expression of pluripotency genes was maintained 48 hours post-SON knockdown (Figure S5G). Junction RT-PCR revealed that SON inhibition led to a significant increase in hTERT-ΔEx2 and a concomitant decrease in full-length telomerase transcripts (Figure 4C and Figure S5F). In agreement with exon-2 skipping leading to compromised hTERT mRNA stability, SON depletion resulted in a significant decrease in TRAP activity in ESCs as well as HeLa cells (Figure 4D-E and S5H-I). In conclusion, our results identify SON as a key regulator of hTERT exon-2 splicing and highlights its role in controlling telomerase activity.

Mutations in SON have been identified in a number of patients diagnosed with intellectual-disability and developmental disorders (41), and one proband was also diagnosed with dyskeratosis congenita (DC), an inherited bone marrow failure syndrome (IBMFS) due to very short telomeres as a result of pathogenic germline variants in the telomerase pathway. DC patients are characterized by the diagnostic triad of nail dystrophy, abnormal skin pigmentation, oral leukoplakia, as well as high rates of bone marrow failure, cancer, pulmonary fibrosis and other complications (42). We identified a female patient (NCI-550-1) with the DC mucocutaneous triad, history of leukopenia, and an additional clinical diagnosis of Zhu-Tokita-Takenouchi-Kim (ZTTK) syndrome (41); an intellectual-disability syndrome associated with cerebral malformations, epilepsy, vision abnormalities, and dysmorphology. The patient is heterozygous for a premature termination allele of the SON gene SON Q96X (chr21:34,921,823, c.286C>T) (Figure 4F). We measured telomere length in granulocytes and lymphocytes using flow cytometry in-situ hybridization (Flow-FISH) and noted very short telomeres (≤1st percentile) in the proband compared to the healthy population (Figure S6). Furthermore, we performed TRAP assay on lysates obtained from peripheral blood mononuclear cells (PBMCs) obtained from the proband and noted substantially reduced telomerase activity relative to a healthy subject (Figure 4G-H). Collectively, our results reveal that SON alternations compromises telomerase activity leading to telomere shortening that might contribute to disease manifestation.

It is well-established that high telomerase levels in the early stages of development resets telomere reserves which is critical for tissue renewal, prevention of degenerative disorders, and possibly setting longevity. Yet, over 30 years since the discovery of telomerase, the fundamental question of how hTERT becomes activated in the inner cell mass and subsequently repressed in differentiated cells remains elusive. Based on our data, we propose a model where hTERT is dually regulated by transcriptional as well as post-transcriptional processes. We show that transcriptional regulation is driven by *cis*-regulatory elements that control hTERT promoter activity. Importantly, our findings indicate that alternative splicing an molecular gatekeeper for hTERT regulation as a function of pluripotency. Specifically, we highlight a previously unanticipated alternative-splice switch centered around exon-2 that ensures the accumulation of full-length hTERT in pluripotent cells. Exclusion of hTERT exon-2 serves to completely repress telomerase activity in somatic cells and enforces the tumor suppressor function of telomere shortening. This is supported by the identification of TERT alternative splice variants in additional species, as evolutionarily distant as Planarians (26, 43, 44). Our study also implicates SON in the regulation of this key alternative splicing switch and we link SON mutations with telomere shortening in a DC patient. At this stage, we cannot exclude that SON mutations impair telomerase activity indirectly by altering levels of other cellular RNAs. However, given the direct role for SON in regulating hTERT exon choice (Figure 4B-C), we favor the scenario where disruption of SON directly impacts telomerase biogenesis. In addition, our results underscore a potential therapeutic benefit for targeting hTERT alternative splicing to block telomerase in cancer cells. Alternatively, enhanced inclusion of hTERT exon-2 might boost telomerase activity in syndromes associated with short telomeres and in regenerative medicine therapies.

## Acknowledgements

We thank Marion Pouillard for her assistance in the early stages of this project. We acknowledge Eros-Lazzerini Denchi and members of the Sfeir lab for comments on the manuscript. We thank Luis Batista for sharing methods for hepatocyte differentiation. Lei Bu, Jerry Shay, and Maria Barna are thanked for reagents. We also thank Ashley S. Thompson for assistance with gene and variant curation. We acknowledge the genome technology core (GTC), high throughput biology (HTB) core, Michael Cammer and the microscopy core at NYU School of medicine. This work was supported in part by a grant from the Irma T. Hirschl foundation to A.S. an NIH fellowship to A.P, and NIH grant RM1HG009491 to J.D.B. The authors would like to dedicate this study to the memory of Woodring E. Wright.

## Author Contributions

A.S. and A.P. conceived the experimental design. All experiments were performed by A.P. A.B. assisted with RNAi screen. M.S. in the lab of J.D.B. helped generate the minigene reporters. S.A.S. evaluated clinical data on the patient with the identified SON mutation. A.S. and A.P. wrote the manuscript. All authors discussed the results and commented on the manuscript.

## Author information

Agnel Sfeir is a co-founder, consultant, and shareholder in Repare Therapeutics. J.D.B. is a founder and director of the following: Neochromosome, Inc., the Center of Excellence for Engineering Biology, and CDI Labs, Inc. and serves on the Scientific Advisory Board of the following: Sangamo, Inc., Modern Meadow, Inc., and Sample6, Inc.

Correspondence and requests for materials should be addressed to A.S.

## METHODS

### Cell culture procedures and treatments

*ARPE-19* cells (ATCC^©^ CRL-2302™), HeLa (ATCC® CCL-2), T-REx™-HeLa (Invitrogen™) and BJ cells (ATCC® CRL-2522™) in Dulbecco’s Modified Eagle Medium (DMEM, Corning™) supplemented with 10% fetal bovine serum (FBS, Gibco™), 2 mM L-glutamine (Gibco™), 100 U/ml Penicillin-Streptomycin (Gibco™), and 0.1 mM MEM non-essential amino acids (Gibco™). ARPE-19 and BJ cell lines were immortalized with hTERT and cultured in the same media conditions. Cells were passaged every 48-72 hours and maintained mycoplasma free by using Plasmocin™ (Invivogen) *per* manufacturer indication. *H7* human embryonic stem cells were a kind gift from Lei Bu and cultured in mTeSR™ Plus (STEMCELL™) media supplemented with 100 U/ml Penicillin-Streptomycin (Gibco). ESCs were passaged every 2-3 days using 500 µM EDTA in PBS and recovered for one day in Y-27632 2HCl (5-10 µM, SelleckChem^®^). *293T* cells used for lentiviral packaging were cultured in DMEM supplemented with 10% bovine calf serum (BCS, Gemini), 2 mM L-glutamine (Gibco™), 100 U/ml Penicillin-Streptomycin (Gibco™), 0.1 mM MEM non-essential amino acids (Gibco™). hTERT minigene expression was induced with 2 µM doxycycline (Sigma-Aldrich^®^) for 48 hours following siRNA tranfections.

iPSC reprogramming of BJ fibroblasts was performed using CytoTune™ - iPS Sendai Reprogramming kit (ThermoFisher) as per manufacturer instructions. Fibroblast differentiation from hESCs was performed in differentiation media containing 15% fetal bovine serum (FBS, Gibco™), 2 mM L-glutamine (Gibco™), 100 U/ml Penicillin-Streptomycin (Gibco™), and 0.1 mM MEM non-essential amino acids (Gibco™). Media was changed without passaging every two days for 21 days when fibroblast differentiation was complete. Cells were subsequently passaged every 4-5 days. Hepatocyte differentiation was performed as described in Mallanna et al. (33) with media changes every 2 days. Briefly, ESCs were plated in Geltrex™ (Gibco) coated 6-well plates and treated with RPMI 1640 (Corning) media containing 2% B27 supplement (w/o insulin, Gibco), and additional recombinant factors including Activin A (100ng/ml, Thermo), BMP4 (10ng/ml, Thermo), FGF2 (20ng/ml, Thermo) for two days. Day 3-5 media contained only Activin A in B27-RPMI 1640 media. Day 6-10 media contained 2% B27 supplement with insulin (Gibco) and BMP4 and FGF2 at previous concentrations. Days 11-15 media contained recombinant HGF (20ng/ml, Thermo) in B27+insulin RPMI 1640 media. Days 16-24 media contained recombinant Oncostatin M (20ng/ml, Thermo) in HCM media lacking EGF (Lonza).

For cell treatments, the following compounds were used: 5-fluoro-uracil (50 µM, Sigma-Aldrich^®^), Isoginkgetin (20 µM, Cayman Chemicals^®^).

The patient was a participant the IRB-approved longitudinal cohort study titled Etiologic Investigation of Cancer Susceptibility in Inherited Bone Marrow Failure Syndromes (NCT-00027274). Informed consent was signed by her parents and data were collected through questionnaires and medical record review (46).

### 4C and ATAC-seq genome-wide sequencing and bioinformatic analysis

Library preparation for Circular Chromosome Conformation Capture (4C) was performed as described in van de Werken et al 2012 (13) with the exception of initial fixation and lysis procedure, which was adapted from Miele et al 2009 (47). 1×10^7^ ARPE or H7 ESCs were dissociated and resuspended in 5ml PBS containing 10% BCS. Cells were crosslinked by adding 5ml of 4% formaldehyde in PBS and incubating at room temperature (RT) for 10 minutes with gentle rocking. Reaction was quenched by adding 2.5ml 2.5M glycine, mixing well and incubating at RT for 5 minutes, then on ice for 15 minutes. Cells were pelleted at 800xg for 10 minutes, resuspended in 1ml of cold Lysis Buffer 1 (10mM Tris pH 8.0, 10mM NaCl, 0.2% IGE-PAL, Protease inhibitor (Roche)) and incubated on ice for 15 minutes. Cells were then dounce-homogenized on ice with a B-type pestle twice for 15 minutes with a minute rest on ice in between. Sample was pelleted at 2500xg for 5 minutes and the pellet was washed with 1x Cutsmart buffer (NEB) and pelleted again. Samples were further processed as described in van de Werken 2012 (13) using DpnII primary digestion and Csp6I secondary digestion. Sequencing library was prepared as described using Expand Long Template polymerase (Roche) and Illumina adaptor sequence-containing primers unique to the hTERT fragend listed in Supplementary Table 1. Sequencing was performed by the NYU Genome Technology Core on Illumina HiSeq 4000 sequencer. Sequencing data was analyzed using the 4Cker software package described in (14) using the recommended parameters.

ATAC-seq was performed on 5×10^5^ ARPE and H7 cells in triplicate as described in Buenrostro et al. 2015 (15). Sequencing of libraries was performed by NYU Genome Technology on Illumina HiSeq 2500 sequencer. Alignment was performed as in Buenrostro et al. 2013 (48) to the hg19 human reference genome and peak calling was done using macs2 using parameters described in Corces et al. 2017 (49). IGV was used to visualize sequencing data tracks and bedtools was used for find regions of overlap between ATAC-seq and 4C data.

### Dual-Luciferase assay for hTERT promoter activity

Dual luciferase reporter plasmid pRF-HCV was a kind gift from Maria Barna (50). HCV promoter was replaced with hTERT core promoter sequence amplified from ESC genomic DNA and cloned into pRF using NcoI restriction enzyme sites. Candidate sequences were similarly amplified from genomic DNA and cloned into BbvCI restriction enzyme site upstream of hTERT promoter. Plasmid DNA was purified using Plasmid Plus Midi Kit (Qiagen) or NucleoBond Xtra Midi Kit (Machery Nagel). Reporter plasmid was introduced to H7 and ARPE cells via transient transfection. Cells were lysed 48 hours after transfection and assayed for luciferase activity using the Dual-Luciferase Reporter Assay System (Promega) according to the manufacturer’s instructions using a Flexstation 3 Multi-mode Microplate Reader (Molecular Devices). Each sample was assayed in triplicate.

### Real-Time RT-qPCR

Total RNA was purified with RNAeasy Mini Kit (Qiagen) or NucleoSpin® RNA Clean-up (Macherey-Nagel) following manufacturer instructions. Genomic DNA was eliminated by on-column digestion with DNaseI. A total of 1 µg of RNA was reverse-transcribed using iScript Reverse Transcription Supermix (Biorad) and qPCR (45 cycles) was performed on a Roche LightCycler480. Reactions were run in triplicates with ssoAdvanced SYBR green Supermix (Biorad) in a total volume of 10µl with standard cycling conditions. Relative gene expression was normalized using HPRT or TBP as housekeeping genes and all calculations were performed in Excel. A list of primers is available in Suppl. Table 1.

### RT-qPCR for hTERT exon-exon junctions

Total RNA was purified with RNAeasy Mini Kit (Qiagen) or NucleoSpin® RNA Clean-up (Macherey-Nagel) following manufacturer instructions. Genomic DNA was eliminated by on-column digestion with DNaseI. A total of 10 µg of RNA was reverse-transcribed using Superscript IV Reverse-Transcription kit (Invitrogen) using an hTERT specific probe in exon-4 (CCTGACCTCTGCTTCCGACAG). Primer was annealed to hTERT RNA at 65C for 5 minutes and then incubated at 0C for 5 minutes before addition of other kit reagents. Reaction was then incubated at 60C for 50 minutes and heat inactivated for 10 minutes at 80C. Each reaction was RNAseH (NEB) treated according to manufacturer instructions and cDNA was purified using MinElute Reaction Cleanup Kit (Qiagen). cDNA concentration was determined using Qubit™ ssDNA Assay Kit (Invitrogen™) and then equalized between samples with ddH_2_O. qPCR (50 cycles) was performed on a Roche LightCycler480. Reactions were run in triplicates with PrimeTime Gene Expression Master Mix (IDT) in a total volume of 10µl with standard cycling conditions. hTERT exon1-2 and exon1-3 expression was normalized using hTERT exon3-4 genes and all calculations were performed in Excel. A list of primers is available in Supp. Table 1.

### CRISPR/Cas9 targeting

H7 RE2^-/-^, RE3^-/-^ and RE2^-/-^RE3^-/-^ double knockouts polyclonal populations were generated as previously described (#). Briefly H7 cells were transfected with a total of 2 gRNAs targeting the 500bp region for RE2 (TTCCCTTGCCCGCTAGAGGG and CCCCCAAGGGAATGAAAAAG) or RE3 (GTGTCTGGATGGACCAGCAG and GCAATGGTAACTCAGTGACT) cloned in a modified version of vector pX458, a kind gift from Feng Zhang (Addgene plasmid # 48138), and a plasmid DNA donor containing 500bp of upstream and downstream homology sequence for either RE2 or RE3 flanking a hPGK-driven Puromycin resistance gene surrounded by two LoxP sites. Five days following transfection, transfected cells were treated with puromycin (250ng/ml) and surviving clones were subjected to genotyping PCR using primers included in Supp. Table 1. Homozygous targeted clones were expanded and infected using lentivirally encoded Cre-recombinase with a Hygromycin resistance gene. Following infection, cells were treated with hygromycin (100ug/ml) and genotyping PCR was performed to confirm excision of the puromycin cassette by fragment size using primers in Supp. Table 1. H7 hTERT^Δin/Δin^ ESCs were obtained by delivering a single-stranded 200nt template oligo containing abutting 100nt sequences from hTERT exon-1 and exon-2 and either an RNP complex of a single gRNA targeting hTERT intron-1 (CGGGGGGAACCAGCGACATG), tcrRNA and wild-type Cas9 protein (IDT) or 2 gRNAs targeting hTERT intron-1 (CGCATGTCGCTGGTTCCCCC and CGGGGGGAACCAGCGACATG) cloned into a modified pX458 plasmid described above. Transfected cells were plated at clonal density and individual clones were picked for genotyping approximately one week later. Genotyping was performed by PCR looking for loss of 104bp in PCR amplicon spanning hTERT exons 1-2 (Supp. Table 1). Genomic DNA was extracted using the Quick-DNA Miniprep Kit (Zymo). Genotyping PCR was performed using Failsafe PCR 2x PreMix H (Lucigen) and Taq polymerase (NEB).

### Lentiviral delivery

Cre-recombinase and Vp64 Cas9-Activation constructs (19) were purchased in the pLenti backbone (Addgene 73795, 61425, 61426) and were introduced by 4 lentiviral infections at 12hr intervals in presence of 8 µg/ml polybrene (Sigma-Aldrich^®^) using supernatant from transfected 293T cells. For targeting Vp64-dCas9 to hTERT promoter, two guide RNA sequences were used in combination (CCAGCTCCGCCTCCTCCGCG and CCAGGACCGCGCTTCCCACG). Vp64 antibiotic resistance genes were replaced with fluorescent protein (mCherry, sfGFP, and tagBFP) genes via several cloning strategies to allow for FACS selection of triple-positive cells containing all Cas9-activation components.

### TRAP Assay for telomerase activity

Telomerase Repeat Addition Processivity (TRAP) was performed as described (#). In brief, cells were dissociated and counted on a manual hemocytometer (Fisher Scientific) using Trypan Blue (Corning) to count viable cells. Cells were pelleted and resuspended in CHAPS lysis buffer (Millipore) with Halt Protease+Phosphatase inhibitor (Thermo) and 3µM β-mercaptoethanol at a concentration of 5×10^3^ cells per µl. Lysates were incubated on ice for 30 minutes, vortexed twice during incubation, and then clarified at 12,000g for 20 minutes. Serial dilutions and heat inactivated samples were prepared and 2µl of lysate or dilution was used in each PCR reaction as described in (18). Cycling conditions were as follows: incubate 30 minutes at 30C, boil 5 minutes at 95C, then melt 30 second at 95C, anneal 30 seconds at 59C, extend 1 minute at 72C; 25 cycles were used for ESC samples, 26 cycles for Hela, and 30 cycles for fibroblast and PBMC samples, and in all cases reaction concluded with a final extension for 10 minutes at 72C. Reactions were run on a 10% acrylamide gel (19:1, Fisher Scientific) and imaged on a ChemiDoc MP apparatus (Biorad).

### RNA-Capture sequencing for hTERT transcripts

Total RNA was purified with RNAeasy Mini Kit (Qiagen) following manufacturer instructions. Genomic DNA was eliminated by on-column digestion with DNaseI. A total of 25 µg of RNA was reverse-transcribed using Superscript IV Reverse-Transcription kit (Invitrogen) in multiple PCR-tubes using a mix of hTERT specific primers in exon-4, exon-9, exon-12, and exon-16 (Supp. Table 1). Primers were annealed to hTERT RNA at 65C for 5 minutes and then incubated at 0C for 5 minutes before addition of other kit reagents. Reaction was then incubated at 60C for 50 minutes and heat inactivated for 10 minutes at 80C. Each reaction was RNAseH (NEB) treated according to manufacturer instructions and second-strand cDNA was synthesized using Second Strand DNA Synthesis kit (NEB) per manufacturer’s instructions. NYU Genome Technology core designed custom probe-library tiling hTERT exons and flanking intragenic sequences using X-Gen probe design software proceeded with Illumina sequencing library preparation of double-stranded cDNA and hybridization and purification of target sequences. Libraries were sequenced using the Illumina MiSeq and data was aligned to hg19 reference genome using Bowtie2 and alternative splicing analysis performed using TopHat.

### Transient Transfection of plasmid DNA and siRNA

Purified plasmid DNA (2-3μg) was introduced to ARPE cells via transient transfection using Lipofectamine 3000 transfection reagent (ThermoFisher) or to H7 ESCs using Genejuice transfection reagent (MilliporeSigma) according to manufacturer’s instructions.

For candidate validation experiments, 2-10 pmol of 4-oligo ON-TARGETplus siRNA pools (Horizon, Dharmacon) or non-targeting control pools (Horizon, Dharmacon) were introduced to H7 ESCs and HeLa cells using 4D-Nucleofector™ (Lonza™) electroporation as per manufacturer instructions for each cell line (ESC: CA-137 in P3 solution, HeLa: CN-113 in SE solution). Splice-blocking morpholino (ASO) for hTERT exon-2 5’ splice site (AGGACACCTGCGGGGGAAGCG) was ordered from Gene Tools, LLC according to their design recommendations with a 3’-Carboxyfluorescein residue to assess transfection efficiency. ASO was delivered to cells by nucleofection as described above.

### hTERT Exon 1-3 splicing minigene assembly by homologous recombination

DNA inserts were PCR amplified with oligos containing the corresponding VEGAS adapters and gel purified while the VEGAS backbone (36) was digested with *Bsa*I and gel purified. ∼100 ng of each fragment along with the linearized VEGAS backbone were transformed using the standard lithium acetate method into *Saccharomyces cerevisiae* strain BY4741 (51) and plated onto SC– Ura plates. After 48 hours colonies were replica plated onto SC–Ura plates containing G418. After an additional two days of growth colonies were screened using PCR to verify the presence of each fragment-fragment junction as well as the initial and terminal backbone-fragment junctions. Colonies containing all junctions were grown overnight in liquid SC–Ura media containing G418. Plasmids were extracted from yeast as follows. 1.5 mL of overnight culture was spun down and resuspended in 250 µL P1 buffer with RNAse (Qiagen) and 200 µL of glass beads (Sigma) in an eppendorf tube and shaken for 10 minutes to break the cells. 250 µL of P2 buffer (Qiagen) was added, mixed by inversion, and incubated for 5 minutes at room temperature. 350 µL of P3 buffer (Qiagen) was added and mixed by inversion. This mixture was spun at 10000xg for 10 minutes in a tabletop centrifuge. The supernatant was transferred to a Zyppy miniprep column (Zymo Research) and spun at 10000xg for one minute; the flowthrough was discarded. The column was washed with 200 µL endo wash buffer, spun at 10000xg for one minute, then washed with 400 µL Zyppy wash buffer and spun again for one minute with flow through discarded. The column was spun one more time to remove residual wash buffer. DNA was eluted with 10 µL of elution buffer. 3 µL of eluted DNA was transformed into *E. coli*. Plasmid DNA recovered from E. coli was digested with AflIII and NotI to release minigene fragment from shuttle vector and clone into pcDNA™5/FRT/TO (Thermo) mammalian expression vector for integration into Hela cells by co-transfection with pOG44-Flpase (Thermo). Hygromycin (150ug/ml) was used to select for cells with integration and Nano-Glo Dual-luciferase assay kit (Promega) was used to confirm expression of both Firefly and Nano-luciferase in clonally isolated cells.

### RNAi Luciferase Screen for hTERT alternative splicing factors

4.5 pmol of Ambion^®^ Silencer^®^ Select siRNA pools (ThermoFisher) was spotted in a 96-well plate format and transfection complexes were formed in OptiMEM (Gibco) using Lipofectamine RNAiMAX (Invitrogen™) according to manufacturer’s instructions. We then introduced the T-REx™-HeLa cell line with heterozygous integration of two minigene constructs suspended in doxycycline-containing DMEM media and incubated for 48 hours at 37C. Cells were lysed and assayed for luciferase activity using the Nano-Glo Dual-Luciferase Reporter Assay System (Promega) according to the manufacturer’s instructions using a EnVision® Multilabel Plate Reader (PerkinElmer). Each plate was assayed in triplicate.

### Absolute quantification of hTERT mRNA

Total RNA was purified with RNAeasy Mini Kit (Qiagen) following manufacturer instructions. Genomic DNA was eliminated by on-column digestion with DNaseI. A total of 1 µg of RNA was hybridized for 24 hours with a custom library of multiply-labelled fluorescent oligos per manufacturer’s instructions to detect specific hTERT exon-exon junctions to derive and absolute quantification of mature spliced transcripts. Predesigned and validated probes against housekeeping genes, TBP and HPRT, were used for subsequent normalization. Probe sequences are listed in Supp. Table 1. Hybridized RNA:probe mixture was then purified and immobilized on nCounter^®^ chips using nCounter Prep Station (NanoString) and then data was acquired using the nCounter^®^ Digital Analyzer (nCounter^®^ FLEX Analysis System, NanoString). All instruments were run and maintained at the NYU Genome Technology Core. Data was analyzed using nSolver™ software package (v4.0, NanoString).

### Immunofluorescence (IF) and microscopy

Cells were plated on 12 mm circular glass coverslips (Fisher Scientific) and analyzed for IF with standard techniques. Briefly, cells were fixed with 4% (v/v) paraformaldehyde in PBS (Santa Cruz Biotechnology, Inc.) for 5 minutes at room temperature. Cells were washed with PBS, permeabilized with 0.5% (v/v) Triton X for 10 minutes and blocked for 30 minutes with PBS containing 3% goat serum (Sigma-Aldrich^®^), 1 mg/ml bovine serum albumin (BSA, Sigma-Aldrich^®^), 0.1% Triton X-100 and 1mM EDTA. Cells were incubated with the same buffer containing primary antibodies for 2 hours at room temperature followed by secondary antibodies incubations for 1 hour at room temperature. Cells were mounted with ProLong Gold Antifade (Thermo Fisher Scientific), imaged on a Nikon Eclipse 55i upright fluorescence microscope at 20X and analyzed with Nikon software. Additional contrast/brightness enhancement and export were performed with Fiji-ImageJ software (52). DNA was counterstained with 5 µg/mL DAPI as needed. A complete list of antibodies used in the study and relative dilutions is available in Suppl. Table 2.

### Single-molecule hTERT mRNA FISH

Cells were plated on 12 mm circular glass coverslips (Fisher Scientific) and analyzed for smiRNA-FISH using techniques described by Tsanov et al. (32). In brief, cells were fixed with 4% (v/v) EM-grade paraformaldehyde (Electron Microscopy Sciences) for 20 minutes at room temperature. Cells were washed with RNAse-free PBS, and permeabilized in 70% ethanol for 1 hour at 4C. Cells were rehydrated in RNAse-free 1x SSC buffer containing 15% (v/v) formamide (Sigma-Aldrich^®^) for 15 minutes and then hybridization solution was applied to coverslips overnight, containing 1x SSC, 15% (v/v) formamide, BSA (2mg/ml, NEB), dextran sulfate (10%, Sigma-Aldrich^®^), VRC (2mM, NEB), tRNA (0.5mg/ml) and 8pmol of flap-annealed probe oligos for hTERT (IDT) designed using Oligostan (32) software, listed in Supp. Table 1. Flap-annealing of probe oligos was done as in Tsanov et al. (32). After hybridization, cells were washed twice in 1X SSC w/ 15% formamide, and twice with 1x PBS, before DNA was counterstained with 5 µg/mL DAPI. Cells were mounted with ProLong Gold Antifade (Thermo Fisher Scientific), imaged on a Nikon Eclipse Ti2 spinning-disc confocal microscope at 60X and analyzed with Nikon software. Additional contrast/brightness enhancement, quantification of foci and export were performed with Fiji-ImageJ software.

### Western blot analysis

Cells were harvested by trypsinization, lysed in RIPA buffer (25 mM Tris-HCl pH 7.6, 150 mM NaCl, 0.1% SDS, 1% NP-40, 1% sodium deoxycholate) at about 10^4^ cell/μl. After 2 cycles of water bath sonication at medium settings lysates were incubated at 4°C on a rotator for additional 30 minutes. Lysates were clarified by spinning 30 minutes at 14800 rpm, 4°C and supernatant protein concentration was quantified with Enhanced BCA protocol (Thermo Fisher Scientific, Pierce). Equivalent amounts of proteins were separated on an SDS-page (approximately 30 µg) and transferred to a nitrocellulose membrane. Membranes were blocked in 5% milk in TBST (137 mM NaCl, 2.7 mM KCl, 19 mM Tris Base and 0.1% Tween-20). Incubation with primary antibodies was performed overnight at 4°C. Membranes were washed and incubated with HRP conjugated secondary antibodies at 1:5000 dilution, developed with Clarity ECL (Biorad) and acquired with a ChemiDoc MP apparatus (Biorad). Antibodies against GAPDH were used as loading control. A full list of antibodies used in the study and relative dilutions is available in Suppl. Table 2.

**Table 1:**
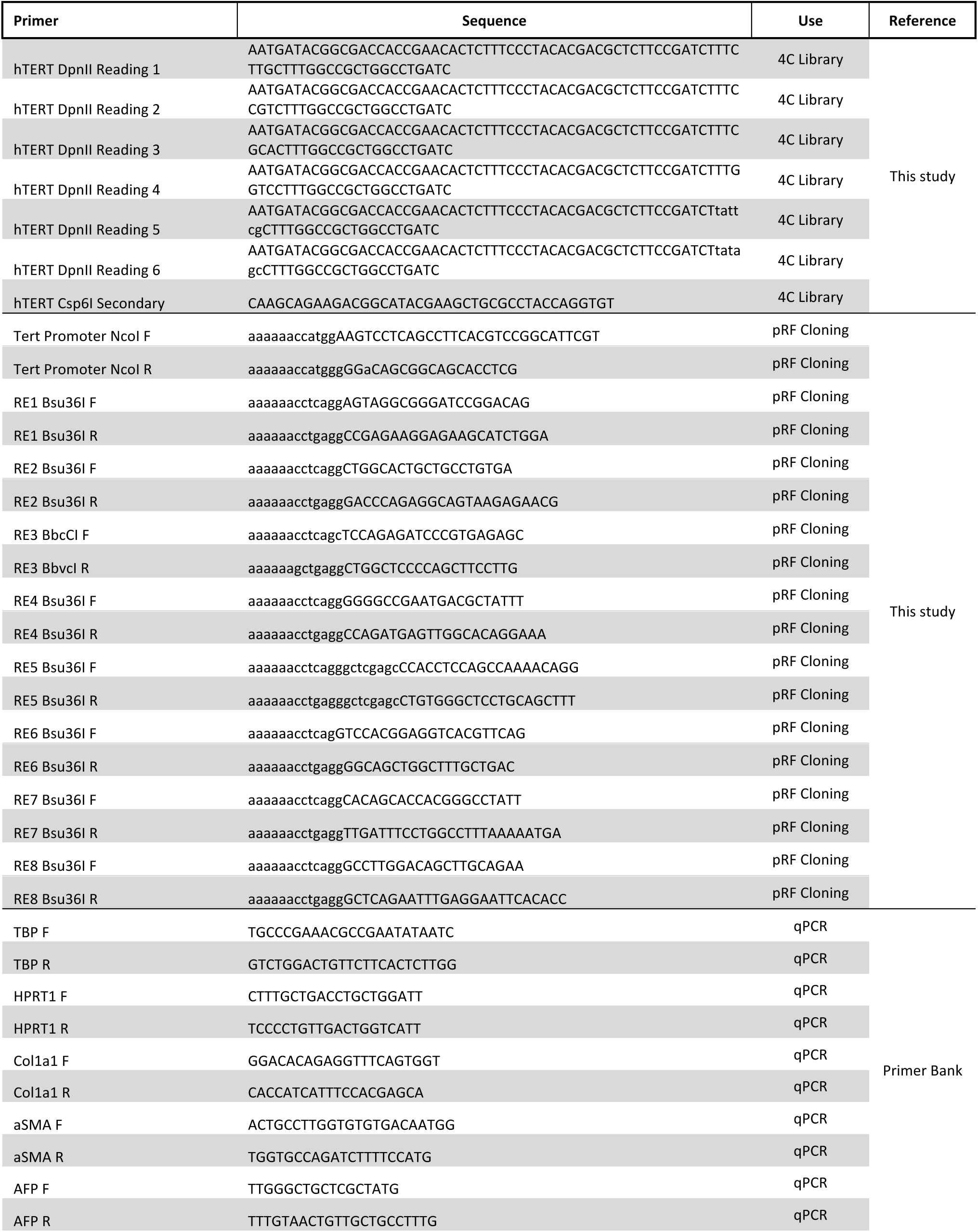
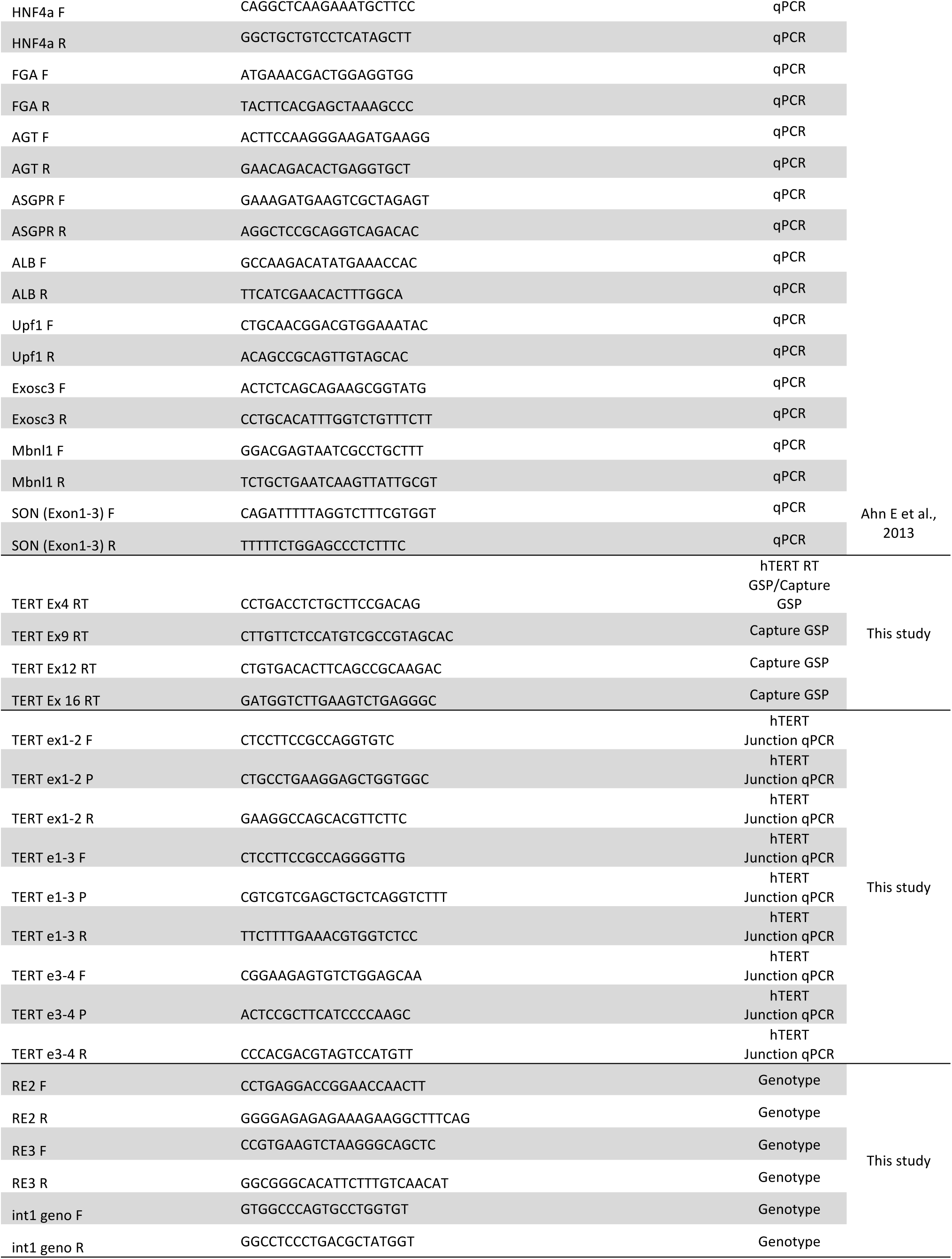
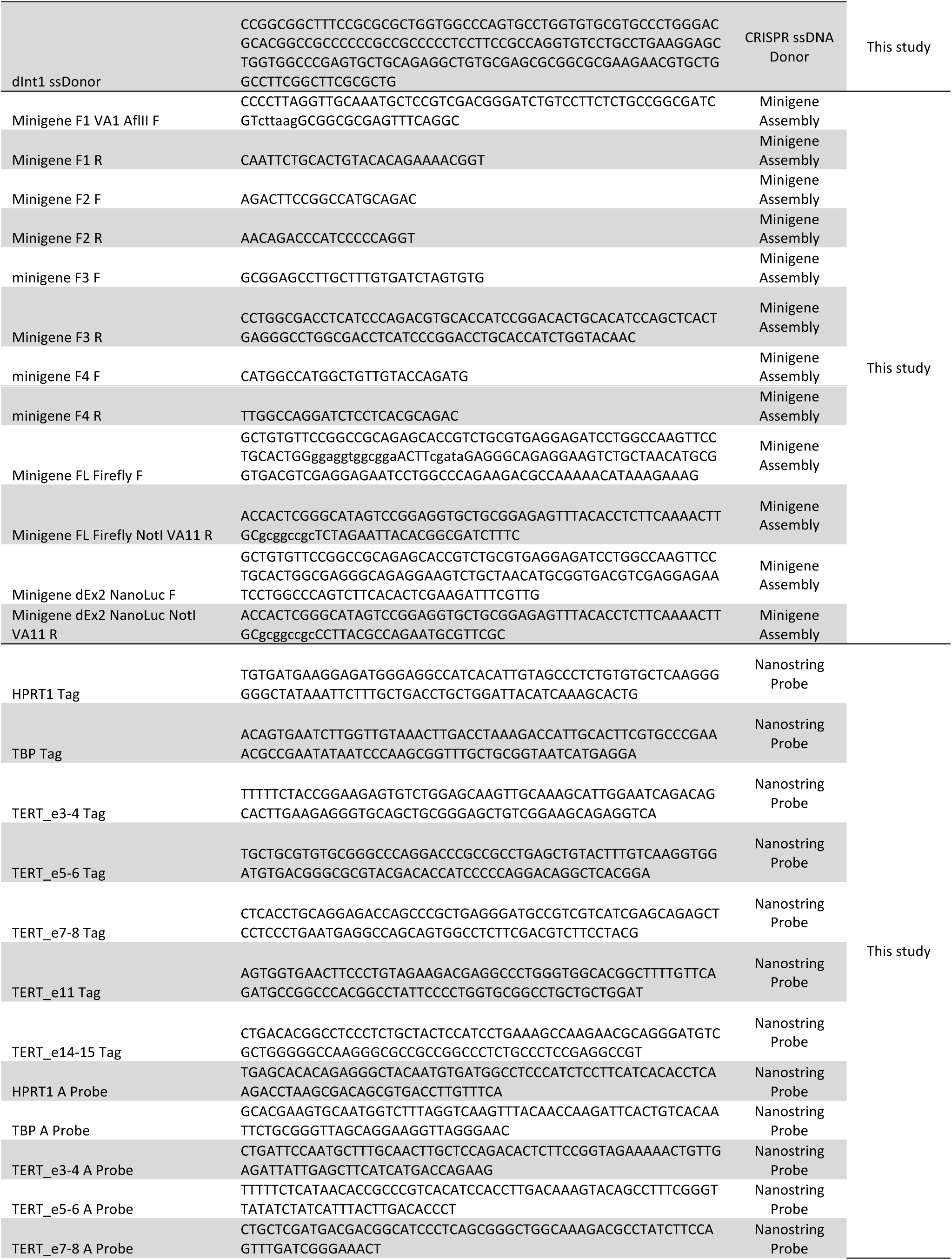
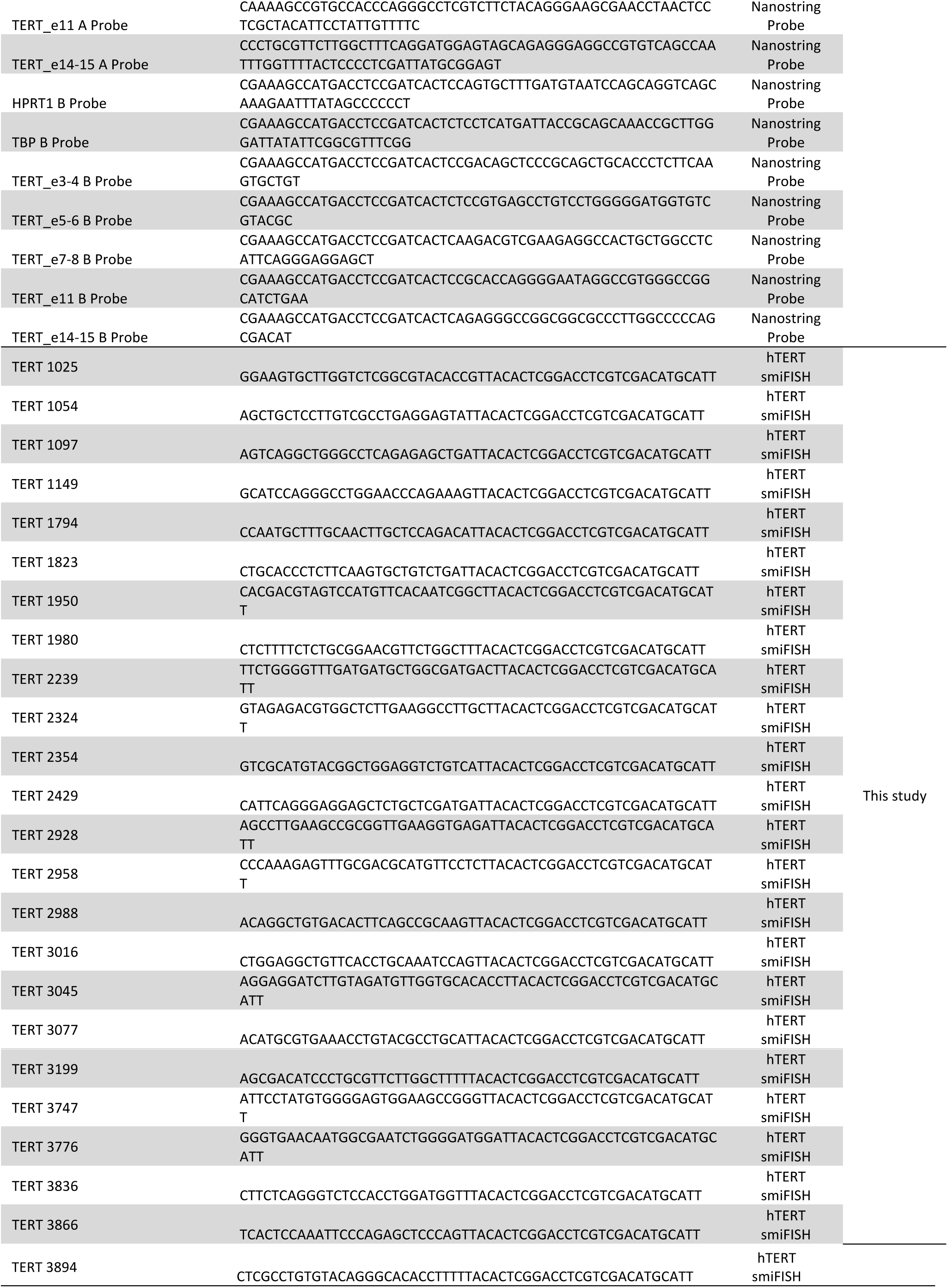
Sequence of oligos used in study.

**Table 3:**
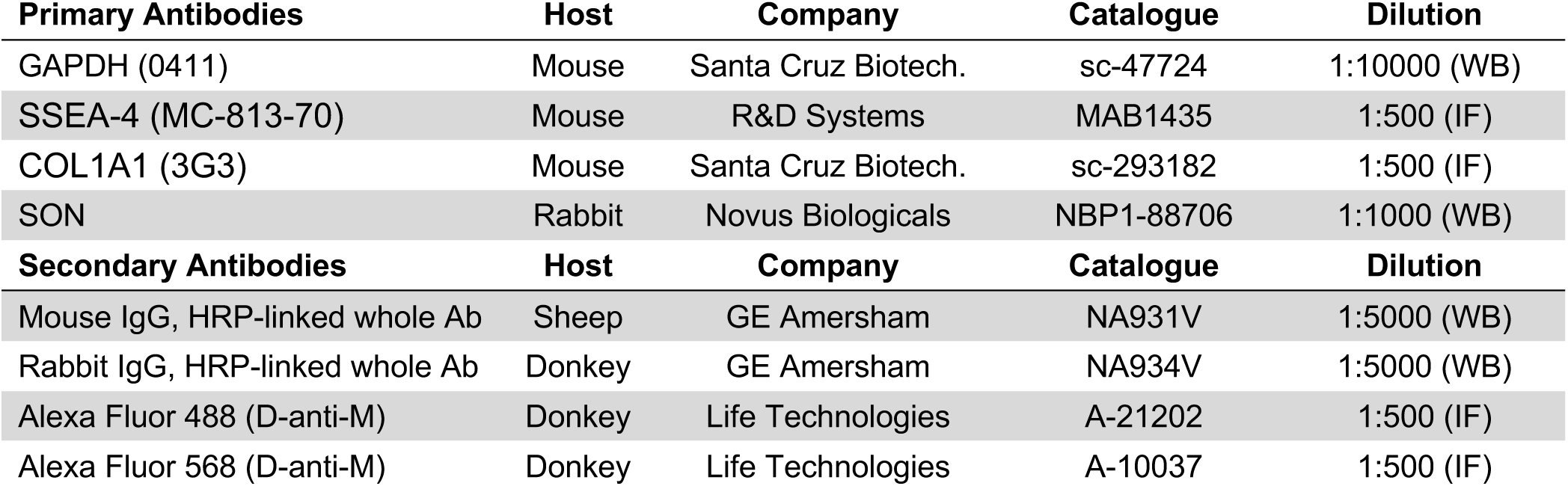
List of Antibodies used throughout the study.

| Primary Antibodies | Host | Company | Catalogue | Dilution |
| --- | --- | --- | --- | --- |
| GAPDH (0411) | Mouse | Santa Cruz Biotech. | sc-47724 | 1:10000 (WB) |
| SSEA-4 (MC-813-70) | Mouse | R&D Systems | MAB1435 | 1:500 (IF) |
| COL1A1 (3G3) | Mouse | Santa Cruz Biotech. | sc-293182 | 1:500 (IF) |
| SON | Rabbit | Novus Biologicals | NBP1-88706 | 1:1000 (WB) |
| Secondary Antibodies | Host | Company | Catalogue | Dilution |
| Mouse IgG, HRP-linked whole Ab | Sheep | GE Amersham | NA931V | 1:5000 (WB) |
| Rabbit IgG, HRP-linked whole Ab | Donkey | GE Amersham | NA934V | 1:5000 (WB) |
| Alexa Fluor 488 (D-anti-M) | Donkey | Life Technologies | A-21202 | 1:500 (IF) |
| Alexa Fluor 568 (D-anti-M) | Donkey | Life Technologies | A-10037 | 1:500 (IF) |

**Table 4:**
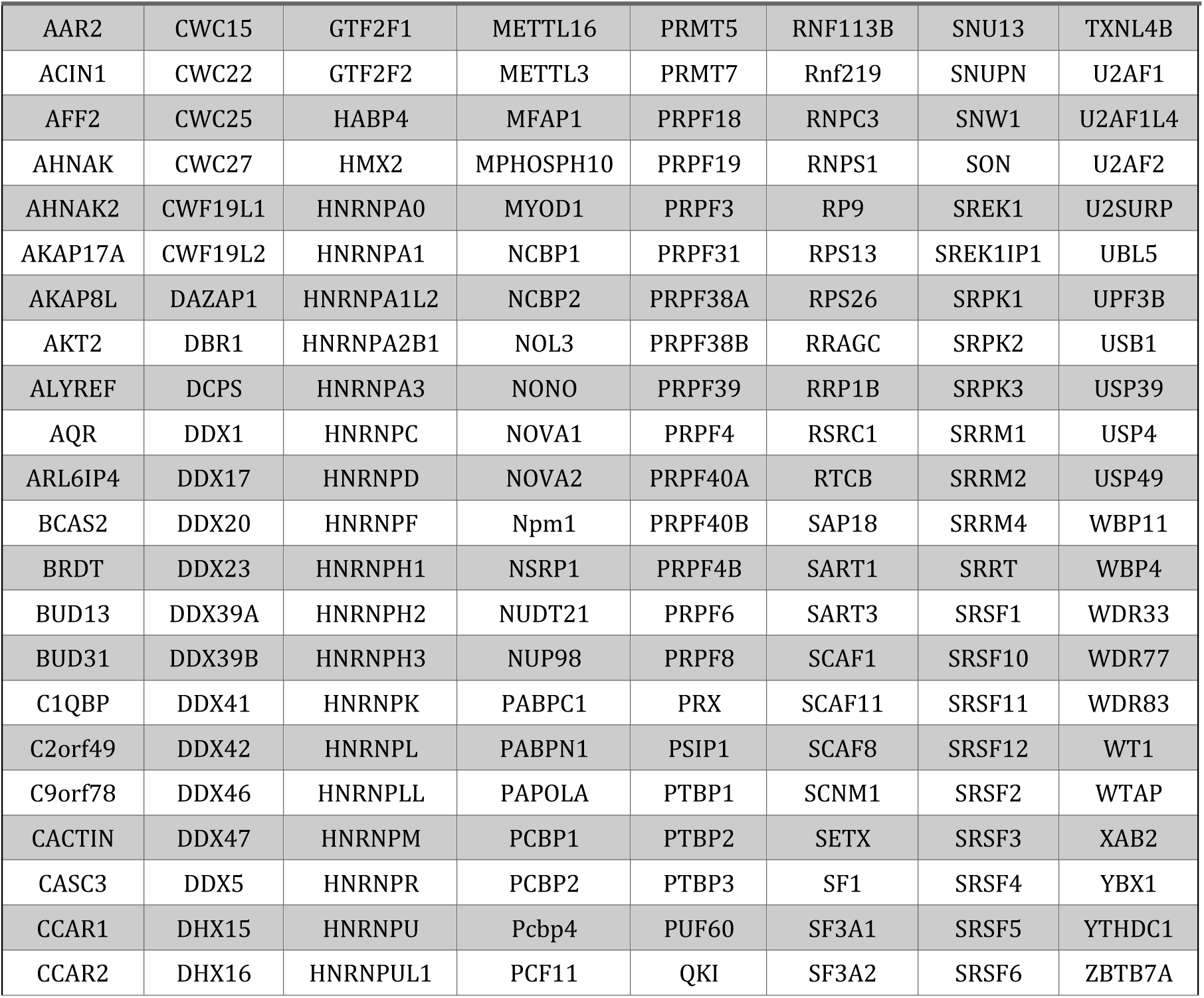
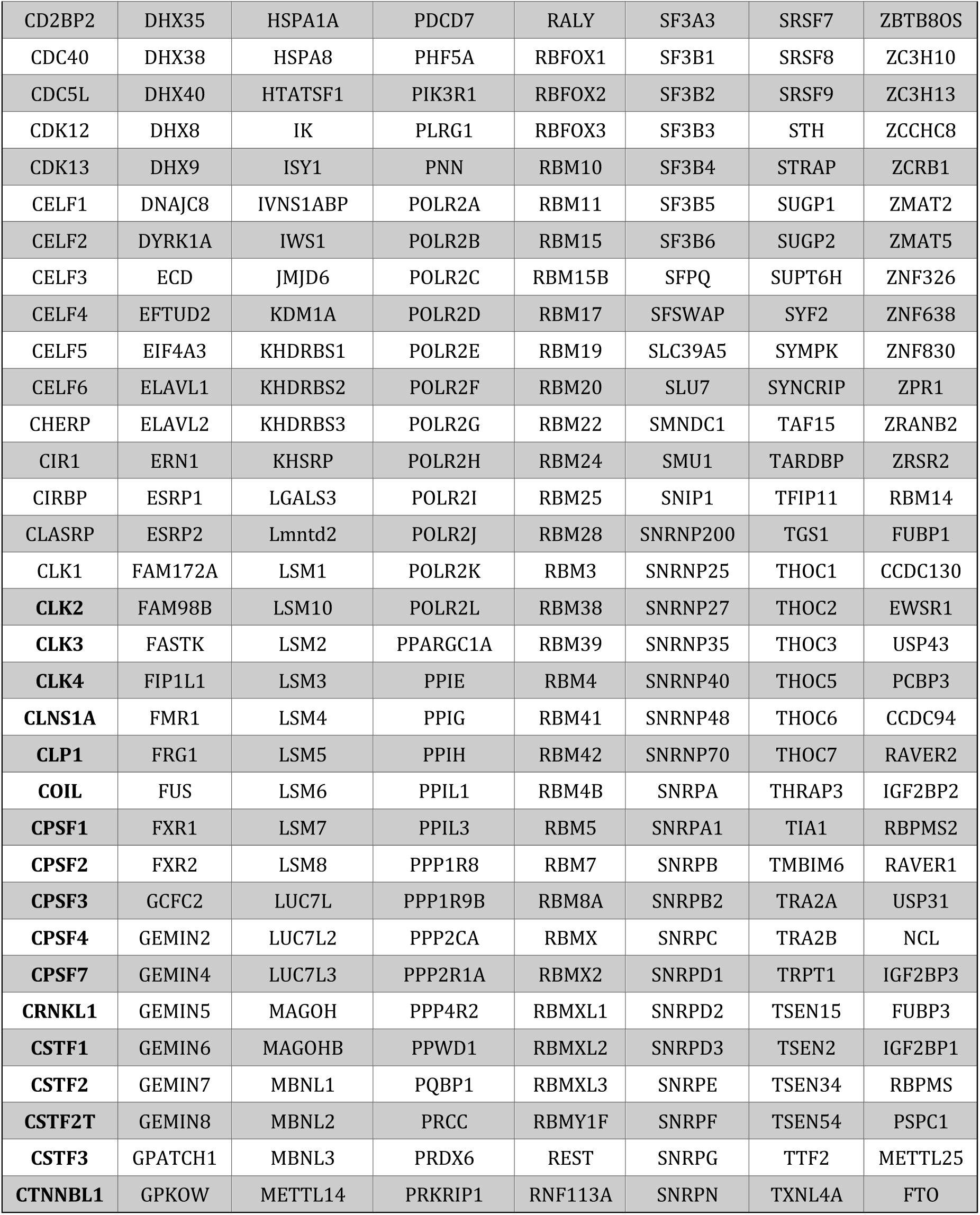
List of RNA-binding proteins and splicing factors tested in RNAi screen.

